# Dissociable roles of dopamine D1 and nicotinic receptors in nicotine-motivated responding and impulsive action in a Go/No-Go self-administration task

**DOI:** 10.64898/2026.07.16.739006

**Authors:** Ranjithkumar Chellian, Guido Huisman, Lara Caglayan, Adriaan W. Bruijnzeel

## Abstract

Cigarette smoking remains one of the most important preventable causes of premature death worldwide. However, the mechanisms sustaining nicotine dependence and relapse are incompletely understood, particularly the role of impulsive action in persistent tobacco use. Dopamine D1 and nicotinic acetylcholine receptors both regulate nicotine-related behavior, but whether they contribute differently to nicotine-motivated responding and nicotine-induced impulsive action is unclear. The present study examined the effects of D1 receptor blockade and stimulation in male and female rats trained to self-administer nicotine intravenously in a Go/No-Go task. Nicotine was available during Go periods but not during No-Go periods, and impulsive action was measured as the percentage of active lever responses during No-Go periods. The D1 receptor antagonist SCH23390 reduced nicotine intake and active lever responding during Go periods and affected the percentage of active lever responses during No-Go periods. However, SCH23390 also reduced inactive lever responding, so its effect on impulsive action could not be separated from a general reduction in operant output. In contrast, the D1 receptor agonist A77636 reduced nicotine intake and Go-period responding without affecting No-Go responding, indicating that D1 receptor stimulation reduced nicotine-motivated responding without altering impulsive action. Rats showed greater No-Go responding during nicotine self-administration than during saline self-administration, indicating that nicotine increased impulsive action rather than general operant responding. The non-selective nicotinic antagonist mecamylamine reduced nicotine-motivated responding and decreased No-Go responding, indicating reduced impulsive action. These findings suggest that nicotinic acetylcholine receptor signaling plays a major role in nicotine-induced impulsive action, whereas D1 receptor signaling contributes more clearly to nicotine-motivated responding. D1 receptor agonism may reduce nicotine-motivated behavior without affecting nicotine-induced impulsive action.

## 1. Introduction

Smoking remains one of the most important preventable causes of disease and premature death worldwide. Tobacco use kills more than 7 million people each year, including approximately 1.6 million nonsmokers exposed to secondhand smoke (WHO 2025a; b). Smoking harms nearly every organ system and is a major cause of cardiovascular disease, stroke, chronic obstructive pulmonary disease, type 2 diabetes, and multiple cancers (Warren et al. 2014). The public health burden is increased by substantial economic costs, including increased healthcare expenditures and lost productivity (Goodchild et al. 2018). Although the adverse health effects of smoking are well established, tobacco use remains difficult to treat because nicotine, the primary addictive constituent of tobacco, produces powerful reinforcing effects and promotes long-lasting neuroadaptations that sustain dependence and relapse (Benowitz 2010; Yamada et al. 2010).

Nicotine is reinforcing, and these effects are at least partly mediated by activation of the mesolimbic dopamine system (Chellian et al. 2023b; Xue et al. 2018). Nicotine stimulates nicotinic acetylcholine receptors (nAChRs) on dopaminergic neurons in the ventral tegmental area (VTA) and thereby increases dopamine release in the nucleus accumbens (NAc) and prefrontal cortex (PFC) (Nisell et al. 1994; Picciotto and Kenny 2021). This dopaminergic response contributes to the acute rewarding effects of nicotine and to the neuroadaptations that promote persistent tobacco use and relapse after abstinence (Corrigall et al. 1992; Koob and Volkow 2010). Consistent with this, pharmacological studies indicate that the mesolimbic dopamine system is critically involved in nicotine reinforcement and nicotine self-administration (Corrigall and Coen 1991; Corrigall and Coen 1994).

A critical but incompletely understood feature of tobacco use disorder is its association with impaired impulse control. Impulsivity comprises several dissociable components, of which impulsive choice and impulsive action are the most relevant to addiction. Impulsive choice refers to the preference for a smaller immediate reward over a larger delayed one, whereas impulsive action refers to the failure to withhold a prepotent motor response (Bari and Robbins 2013; Winstanley 2011). The present study focuses on impulsive action, the form of impulsivity that is most closely tied to deficits in inhibitory control. This form of impulsivity is measured in rodents with tasks such as the Go/No-Go task and the five-choice serial reaction time task (5-CSRTT), which differ in design but both index the failure to withhold a prepotent response, recorded as responding during the No-Go period in the Go/No-Go task and as premature responding in the 5-CSRTT. Studies using these tasks have shown that nicotine increases impulsive action. Acute and repeated nicotine increase premature responding in the 5-CSRTT, an effect reversed by the nAChR antagonist mecamylamine and therefore mediated by nAChR activation (Blondel et al. 2000), and acute nicotine similarly increases impulsive action in a Go/No-Go task (Kolokotroni et al. 2011).

Impulsive behavior has been implicated in the initiation of drug use, the escalation of drug intake, maintenance of dependence, and vulnerability to relapse (Perry and Carroll 2008; Verdejo-García et al. 2008). Prefrontal dopamine signaling plays an important role in top-down inhibitory control, and disruption of this system is associated with impaired response inhibition in people with psychiatric and substance use disorders (Dalley et al. 2011; Robbins et al. 2012). Consistent with this, tobacco smokers show deficits in inhibitory control compared with nonsmokers in Go/No-Go tasks (Luijten et al. 2011; Motka et al. 2025). However, it remains unclear whether impulsive action is primarily a pre-existing vulnerability factor for developing nicotine addiction, a consequence of chronic nicotine use, or both. Preclinical work has begun to disentangle these dynamics. Studies have shown that high baseline impulsive action specifically predicts the rapid acquisition of nicotine self-administration in rats, whereas impulsive choice acts as a stronger predictor of high intake and relapse vulnerability (Diergaarde et al. 2008).

Dopamine D1 receptors are expressed throughout mesocorticolimbic circuits, including the NAc, PFC, and striatum, and are positioned to regulate both drug-motivated behavior and inhibitory control. Dopamine modulation of the PFC is complex, and D1 receptor signaling follows an inverted-U-shaped relationship in which both insufficient and excessive D1 receptor activation can impair cognitive performance (Seamans and Yang 2004; Vijayraghavan et al. 2007; Zahrt et al. 1997). The D1 receptor antagonist SCH23390 reduces nicotine self-administration, which supports a role for D1 receptor signaling in nicotine reinforcement (Corrigall and Coen 1991). More recent work indicates that both D1-like receptor blockade and D1-like receptor stimulation can decrease operant responding for nicotine, although D1-like receptor blockade may also involve broader changes in operant performance or motivation (Chellian et al. 2022; Chellian et al. 2023b).

Dopamine D1 receptor signaling has also been linked directly to nicotine-induced impulsive action. In the 5-CSRTT, nicotine increases impulsive responding without altering attentional accuracy, and this increase is attenuated by SCH23390, implicating D1 receptors in the expression of nicotine-related impulsive action (Balachandran et al. 2018). Despite this evidence linking dopaminergic signaling to nicotine-induced impulsive action, it remains unknown how D1 receptor signaling regulates impulsive action when nicotine is actively self-administered, because previous studies have relied on non-contingent nicotine administration (Balachandran et al. 2018). This is an important gap, as tobacco use disorder is characterized by voluntary, response-contingent drug intake, in which drug-taking and the failure of inhibitory control occur within the same behavioral context. The Go/No-Go task is well suited to model this because it can be implemented under a nicotine self-administration schedule, in which active lever responding is reinforced with intravenous nicotine during Go periods and reinforcement is withheld during No-Go periods. This procedure permits nicotine-motivated responding and impulsive action to be assessed concurrently within a single session under conditions of active drug intake. In the present study, we adapted a Go/No-Go self-administration procedure initially used to study impulsive action during cocaine self-administration to examine how dopamine D1 receptor signaling regulates nicotine-motivated responding and impulsive action in male and female rats (Zapata et al. 2017; Zapata and Lupica 2021). Male and female rats were trained to self-administer nicotine intravenously under a Go/No-Go schedule as in our previous work (Chellian et al. 2026). Nicotine intake and active lever responding during Go periods were used as measures of nicotine-motivated behavior, whereas the percentage of active lever responses during No-Go periods was used as an index of impulsive action. We examined the effects of the D1 receptor antagonist SCH23390 and the D1 receptor agonist A77636 on task performance. In addition, rats self-administering saline were compared with those self-administering nicotine to determine whether the elevated impulsive action requires active nicotine intake or whether chronic drug exposure produces a persistent deficit in inhibitory control. We also tested the non-selective nAChR antagonist mecamylamine to compare dopaminergic and cholinergic contributions to nicotine-motivated responding and impulsive action. Sex was included as a biological variable throughout the study. By dissociating the effects of D1 receptor modulation on Go-period responding from No-Go-period impulsive responding, this study aimed to clarify how dopaminergic signaling regulates reward-driven action and inhibitory control in the context of nicotine self-administration.

## 2. Material and methods

### 2.1. Animals

Adult male (220–280 g, 8-9 weeks of age; N=14) and female (180–225 g, 8-9 weeks of age; N=18) Wistar rats were purchased from Charles River (Raleigh, NC). The rats were housed with a rat of the same sex in a climate-controlled vivarium on a reversed 12 h light-dark cycle (light off at 7 AM). The rats were handled for 2-3 min per day for several days before surgery. Following jugular catheter implantation, rats were singly housed for the remainder of the study. Food and water were available ad libitum in the home cage throughout the study. The experimental protocols were approved by the University of Florida Institutional Animal Care and Use Committee (IACUC). All experiments were performed in accordance with relevant IACUC guidelines and regulations and in compliance with ARRIVE guidelines 2.0 (Animal Research: Reporting of *In Vivo* Experiments).

### 2.2. Drugs

For intravenous self-administration, (-)-nicotine hydrogen tartrate (NIDA Drug Supply Program) was dissolved in sterile saline (0.9 % sodium chloride), and the pH was adjusted to 7.2 ± 0.2 using 1M NaOH. Rats self-administered nicotine at a dose of 0.03 mg/kg/inf (expressed as base) in a volume of 0.1 ml/inf. For drug treatments, (-)-nicotine hydrogen tartrate, SCH23390 hydrochloride (Tocris bioscience, Minneapolis, MN), A77636 hydrochloride (Tocris bioscience, Minneapolis, MN), and mecamylamine hydrochloride (NIDA Drug Supply Program) were dissolved in sterile saline and administered subcutaneously (SC) in a volume of 1 ml/kg body weight. Nicotine doses are expressed as base, while SCH23390, A77636, and mecamylamine doses are expressed as salt.

### 2.3. Experimental design

A schematic overview of the experimental timeline is presented in Figure 1. Rats were surgically implanted with catheters in the jugular vein and allowed a minimum 7-day recovery period. Following recovery, male (N=14) and female (N=18) rats were trained to self-administer nicotine during fifteen 2-h sessions. This was followed by 21 days of nicotine self-administration under a Go/No-Go training paradigm. After completion of the Go/No-Go training sessions, the effects of SCH23390, A77636, and mecamylamine on nicotine self-administration in the Go/No-Go task were evaluated. In addition, a separate comparison examined responding in rats self-administering saline under the Go/No-Go schedule.

**Fig. 1.**
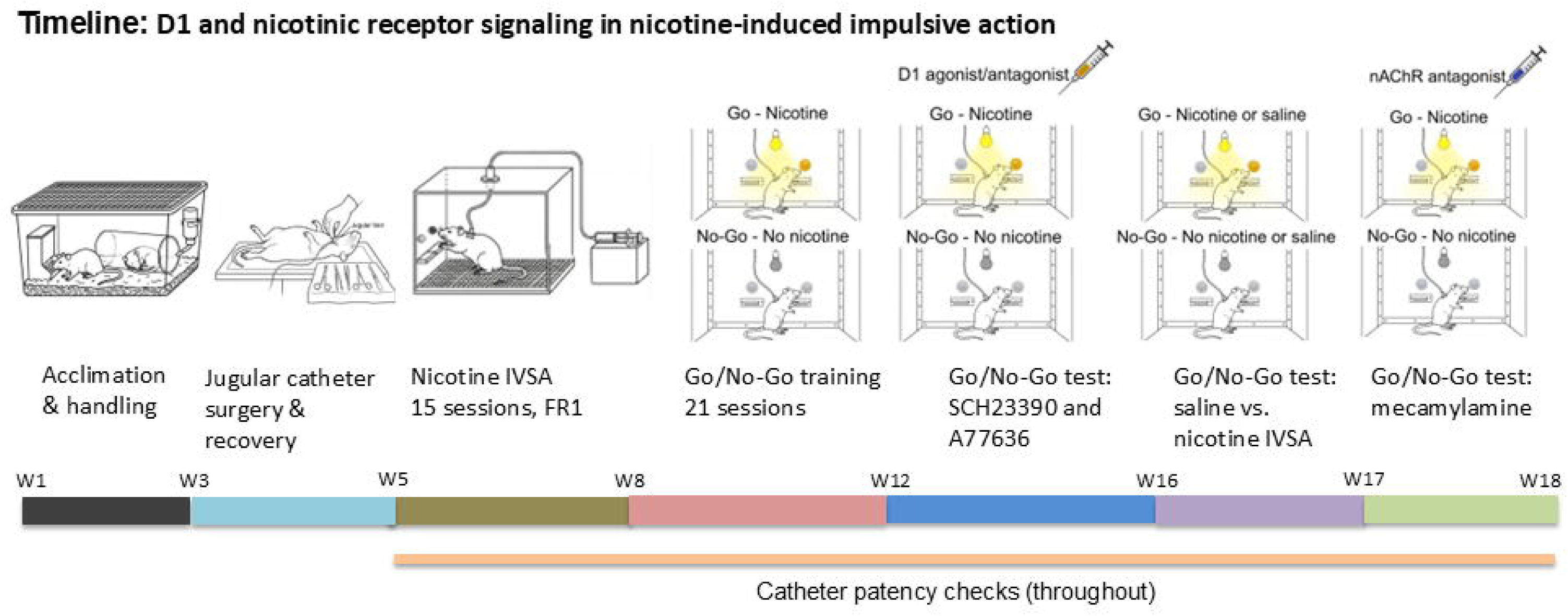
Schematic overview of the experimental timeline for the Go/No-Go nicotine self-administration study. Male and female rats were acclimated and handled before surgical implantation of a jugular catheter, followed by a minimum 7-day recovery period. Rats then acquired intravenous nicotine self-administration during fifteen 2-h sessions under an FR1 schedule. This was followed by 21 sessions of nicotine self-administration under a Go/No-Go training paradigm with alternating Go periods, during which nicotine was available, and No-Go periods, during which nicotine was unavailable. After completion of Go/No-Go training, the effects of the dopamine D1 receptor antagonist SCH23390 and the dopamine D1 receptor agonist A77636 on performance in the Go/No-Go task were assessed. Rats were then tested under saline and nicotine self-administration conditions in the Go/No-Go task. Finally, the effects of the non-selective nAChR antagonist mecamylamine were assessed. Catheter patency was monitored throughout the study. The study began with 14 males and 18 females. The final animal numbers for each experimental phase are reported in the corresponding figure legends.

### 2.4. Intravenous catheter implantation

The catheters were implanted as described before (Chellian et al. 2024a; b). The rats were anesthetized with an isoflurane-oxygen vapor mixture (1-3%) and prepared with a catheter in the right jugular vein. The catheters consisted of polyurethane tubing (length 10 cm, inner diameter 0.64 mm, outer diameter 1.0 mm, model 3Fr, Instech Laboratories, Plymouth Meeting, PA). The right jugular vein was isolated, and the catheter was inserted 2.9 cm for males and 2.5 cm for females. The tubing was then tunneled subcutaneously and connected to a vascular access button (Instech Laboratories, Plymouth Meeting, PA). The button was exteriorized through a 1-cm incision between the scapulae. During the 7-day recovery period, the rats received daily infusions of the antibiotic Gentamycin (4 mg/kg, IV, Sigma-Aldrich, St. Louis, MO). A sterile heparin solution (0.1 ml, 50 U/ml) was flushed through the catheter before and after administering the antibiotic and after nicotine self-administration. After flushing the catheter, 0.05 ml of a sterile heparin/glycerol lock solution (500 U/ml, Instech Laboratories, Plymouth Meeting, PA) was infused into the catheter. The animals received carprofen (5 mg/kg, SC) daily for 72 hours after the surgery.

### 2.5. Acquisition of nicotine self-administration

Following recovery from surgery, rats were trained to acquire nicotine self-administration (0.03 mg/kg/infusion) under a fixed ratio 1 (FR1) schedule with a 10-second time-out (TO) in sound- and light-attenuated operant chambers (Med Associates, St. Albans, VT). Rats self-administered nicotine during daily 2 h sessions for 15 sessions. Nicotine self-administration sessions were conducted five days per week for three weeks. Responding on the active lever resulted in the delivery of a nicotine infusion (0.1 ml infused over a 6.5-s period). The infusion was paired with a cue light above the active lever, which remained illuminated throughout the time-out period. Responding on the inactive lever was recorded but did not have scheduled consequences. Both levers were retracted during the time-out (TO) period.

### 2.6. Go/No-Go nicotine self-administration task

The Go/No-Go self-administration task was conducted as previously described by Zapata et al. using cocaine (Zapata et al. 2017). Following the acquisition of nicotine self-administration, rats were trained to self-administer nicotine under the Go/No-Go schedule. Each session lasted 2 h and consisted of six alternating 20-minute intervals: three Go periods (nicotine available; 0–20 min, 40–60 min, and 80–100 min) and three No-Go periods (nicotine unavailable; 20–40 min, 60–80 min, and 100–120 min). Sessions always began with a Go period for all rats. Nicotine availability during the Go periods was signaled by illumination of the house light. During these periods, responding on the active lever resulted in an infusion of nicotine (0.03 mg/kg/infusion; 0.1 ml infused over 6.5 seconds) under a fixed ratio 5 (FR5) schedule with a 10-s timeout (TO). Each infusion was paired with a cue light above the active lever, which remained illuminated throughout the timeout period. The house light was turned off during the timeout period and turned on afterward to signal continued nicotine availability. Responses on the inactive lever were recorded but had no programmed consequences. Both levers were retracted during the timeout period. During the No-Go periods, the house light remained off to indicate that nicotine was unavailable. Lever presses were recorded during this period, but neither the active nor inactive lever produced any scheduled consequences. We assessed impulsive action using this task, specifically measuring the failure to inhibit responding during the No-Go period (commission errors). This task is a well-established paradigm for evaluating motor inhibition (Bari and Robbins 2013). The house light acts as a discriminative stimulus for nicotine availability during Go periods, and impulsive action is therefore defined as the inability to withhold responding when the learned No-Go signal indicates that nicotine is unavailable. Impulsive responding was quantified as the percentage of active lever responses emitted during No-Go periods relative to total active lever responses across both Go and No-Go periods, calculated as: [No-Go active lever responses / (Go active lever responses + No-Go active lever responses) × 100](Zapata and Lupica 2021). This measure indexes commission errors, reflecting impaired inhibitory control when nicotine was unavailable. Because nicotine infusions during Go periods were obtained under an FR5 schedule, overall active lever responding was higher during Go intervals. Consequently, relying on raw counts of No-Go responses may misrepresent impulsivity, as such differences can arise from variations in overall response output or reinforcement contingencies rather than true deficits in inhibitory control. Expressing commission errors as a proportion of total active-lever responses accounts for these differences and provides a rate-independent measure of inhibitory control. Go/No-Go training was conducted for 21 sessions, five days per week.

### 2.7. Effects of SCH23390 and A77636 on the Go/No-Go nicotine self-administration task

Following the Go/No-Go training sessions, the effects of SCH23390 and A77636 were assessed on nicotine self-administration behavior within the Go/No-Go task. Each drug was administered 15 minutes prior to the Go/No-Go nicotine self-administration (0.03 mg/kg/infusion) session. SCH23390 (0, 0.003, 0.01, and 0.03 mg/kg, SC), and A77636 (0, 0.1, and 0.3 mg/kg, SC) were administered using a Latin square design. The highest dose of A77636, 1 mg/kg, was not included in the Latin square design and was administered last. The doses of SCH23390 and A77636 were based on our previous studies of nicotine self-administration in rats (Chellian et al. 2022; Chellian et al. 2023a). There was minimum 48-hour interval between successive test doses of SCH23390 or A77636. Daily Go/No-Go nicotine self-administration sessions were conducted between the treatment sessions. The treatments with A77636 began 72 hours after the final SCH23390 session. Go/No-Go nicotine self-administration sessions were conducted five days per week, and at least one baseline Go/No-Go nicotine self-administration session was conducted before each Latin square dose of SCH23390 or A77636.

### 2.8. Comparison of nicotine and saline self-administration in the Go/No-Go task

After completion of A77636 treatment experiments, a separate comparison was conducted to evaluate responding for saline versus nicotine under the Go/No-Go schedule. Rats continued to undergo daily Go/No Go nicotine self administration, and ninety six hours after the final drug treatment session, they were assigned to a within subject crossover design. During the first test phase, half of the animals self-administered saline, while the remaining animals self-administered nicotine (0.03 mg/kg/infusion) under identical Go/No-Go task conditions. Following this initial test, animals underwent two standard Go/No-Go nicotine self-administration sessions to re-establish baseline responding before crossing over to the alternate condition. Animals that initially self-administered saline subsequently self-administered nicotine, and animals that initially self-administered nicotine subsequently self-administered saline. Go/No-Go nicotine self-administration sessions were conducted five days per week, and at least one baseline Go/No-Go nicotine self-administration session was conducted before each crossover IVSA session.

### 2.9. Effects of mecamylamine on the Go/No-Go nicotine self-administration task

Following the saline and nicotine self administration Go/No Go sessions, the effects of mecamylamine on nicotine self administration behavior were assessed using the Go/No Go task. Ninety six hours after the final saline and nicotine self administration Go/No Go sessions, mecamylamine (0 and 2 mg/kg, SC) was administered using a Latin square design. Mecamylamine was tested at a single dose, which was based on our dose-response characterization of mecamylamine in the same Go/No-Go nicotine self-administration procedure (Chellian et al. 2026). Mecamylamine was administered 15 minutes prior to the Go/No Go nicotine self administration (0.03 mg/kg/infusion) session. There was a minimum 48 hour interval between successive test doses of mecamylamine. Daily Go/No Go nicotine self administration sessions were conducted between treatment sessions. Go/No Go nicotine self administration sessions were conducted five days per week, and at least one baseline Go/No Go nicotine self administration session was conducted before each Latin square dose of mecamylamine.

### 2.10. Catheter patency test

Catheter patency was assessed during the self-administration period by administering 0.2 ml of the ultra-short-acting barbiturate Brevital (1% methohexital sodium). A rapid loss of muscle tone was considered indicative of a patent catheter. Rats that did not exhibit this response were excluded from subsequent analyses (See Table S1). One male and four female rats failed the Brevital test during the acquisition period and were excluded from further analyses. Three male and three female rats failed the Brevital test during the Go/No-Go training period and were excluded. Additionally, one female rat was excluded during the SCH23390 treatment phase and one female rat was excluded during the A77636 treatment phase because they did not respond to Brevital.

### 2.11. Statistics

Data were analyzed with SPSS Statistics version 31 and GraphPad Prism version 11. The figures were generated using GraphPad Prism version 11. Lever pressing, percentage of active lever responses, nicotine intake, and infusion latency data were analyzed using two-way or three-way analysis of variance (ANOVA) with repeated measures where appropriate. The factors included treatment, sex, session, and lever (active vs. inactive). Significant interactions and main effects were further examined using the Bonferroni post hoc test. The alpha level for statistical significance was set at p < 0.05.

## 3. Results

### 3.1. Acquisition of nicotine self-administration

Male and female rats acquired nicotine self-administration over 15 sessions. During this period, both male and female rats responded more on the active lever than the inactive lever, with no effect of sex (Fig. S1A, Lever F1,25=8.151, P < 0.01; Sex F1,25=1.17, NS; Lever × Sex F1,25=0.562, NS). Active lever responding increased over time during the early acquisition sessions and then stabilized in both males and females, whereas inactive lever responding showed no systematic change across sessions (Fig. S1A, Session F14,350=3.461, P < 0.001; Session × Sex F14,350=0.739, NS; Lever × Session F14,350=4.47, P < 0.001; Lever × Session × Sex F14,350=1.307, NS). Nicotine intake initially increased during the initial acquisition sessions and subsequently stabilized in males and females (Fig. S1B, Session F14,350=15.416, P < 0.001; Session × Sex F14,350=1.41, NS; Sex F1,25=1.64, NS).

### 3.2. Go/No-Go training

After the acquisition phase, male and female rats were trained to self-administer nicotine in a Go/No-Go task for 21 sessions.

#### Go period

During the Go period, active lever responses were higher than inactive lever responses, with no effect of sex (Fig. 2A, Lever F1,19=167.622, P < 0.001; Sex F1,19=0.034, NS; Lever × Sex F1,19=1.805, NS). Active lever responding increased during the initial sessions and then remained relatively stable in both males and females, while inactive lever responses were relatively stable across sessions (Fig. 2A, Session F20,380=4.053, P < 0.001; Session × Sex F20,380=1.42, NS; Lever × Session F20,380=4.66, P < 0.001; Lever × Session × Sex F20,380=0.782, NS). Nicotine intake increased during the initial sessions and then remained relatively stable in both males and females (Fig. 2B, Session F20,380=5.116, P < 0.001; Session × Sex F20,380=1.222, NS; Sex F1,19=0.376, NS).

**Fig. 2.**
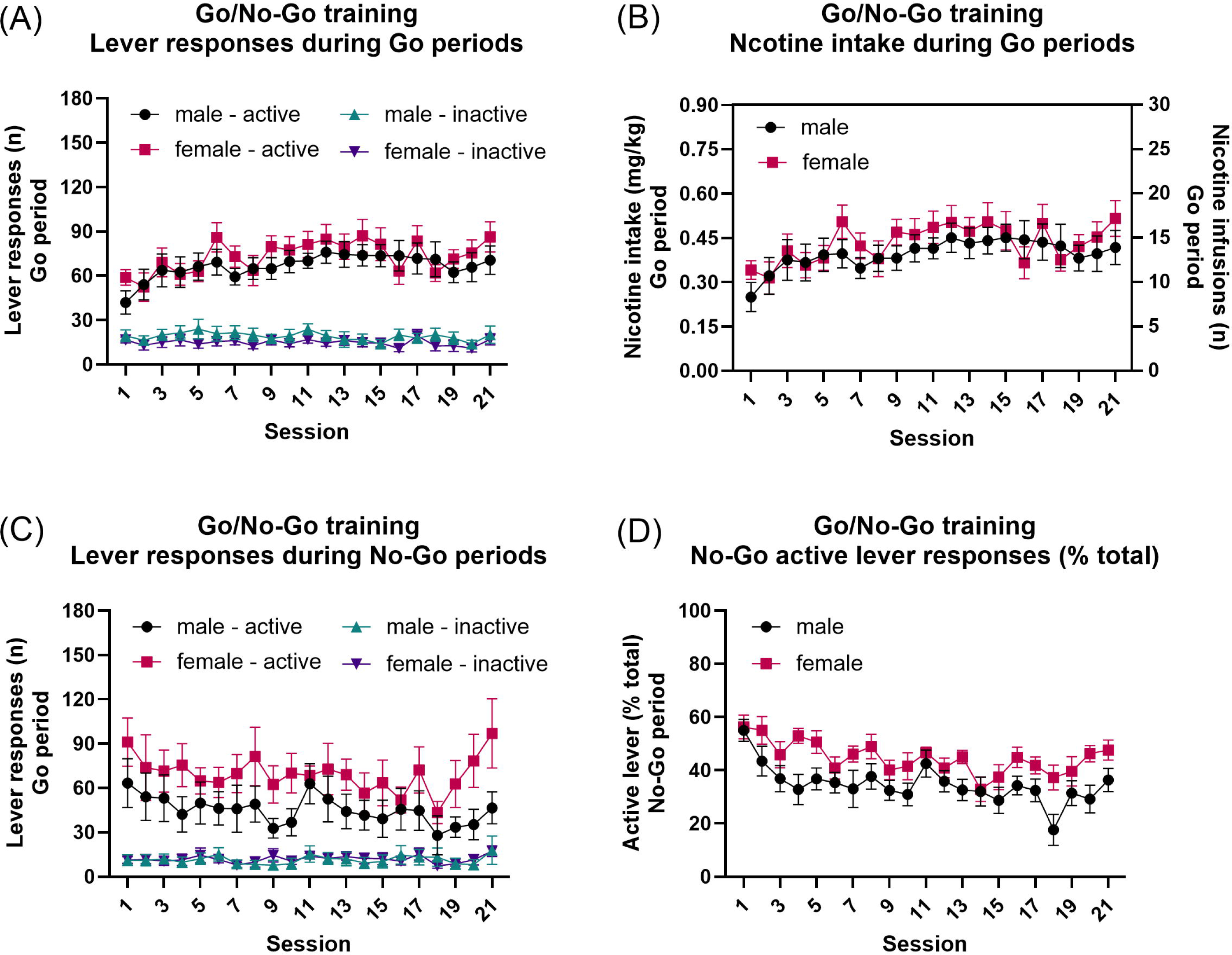
Nicotine self-administration and responding during Go/No-Go training. (A) Active and inactive lever responses during the Go periods across the 21 Go/No-Go training sessions. Rats responded more on the active lever than the inactive lever, and active lever responding increased during the initial sessions and then stabilized. (B) Nicotine intake during Go periods increased during the initial sessions and then remained relatively stable. (C) Active and inactive lever responses during the No-Go periods. Rats responded more on the active lever than the inactive lever during No-Go periods, although active lever responding changed across training. (D) The percentage of active lever responses during the No-Go periods decreased across the initial training sessions and then stabilized, consistent with the acquisition of inhibitory control. Females showed a higher percentage of active lever responses during No-Go periods than males. Males, N=10; females, N=11. Data are expressed as mean ± SEM.

#### No-Go period

During the No-Go period, the rats responded more on the active lever than the inactive lever, with no effect of sex (Fig. 2C, Lever F1,19=56.607, P < 0.001; Sex F1,19=1.738, NS; Lever × Sex F1,19=3.899, NS). Lever responding varied across sessions over time, driven primarily by changes in active lever responding, while inactive lever responding remained relatively stable over time, with no effect of sex (Session F20,380=2.004, P < 0.01; Session × Sex F20,380=0.786, NS; Lever × Session F20,380=1.949, P < 0.01; Lever × Session × Sex F20,380=0.654, NS). The percentage of active lever responses during the No-Go period decreased from the initial sessions and then remained relatively stable over time, with a significant effect of sex (Fig. 2D, Session F20,380=5.557, P < 0.001; Sex F1,19=6.727, P < 0.05; Session × Sex F20,380=1.031, NS).

### 3.3. Go/No-Go task: SCH23390 treatment

#### Go period

SCH23390 treatment decreased active and inactive lever responses, as well as nicotine intake during the Go period, with females showing higher overall active lever responding and nicotine intake than males (Fig. 3A, Active lever: Treatment F3,54=45.787, P < 0.001; Treatment × Sex F3,54=0.768, NS; Sex F1,18=5.117, P < 0.05; Fig. 3B, Nicotine intake: Treatment F3,54=44.378, P < 0.001; Treatment × Sex F3,54=0.82, NS; Sex F1,18=4.995, P < 0.05; Fig. 3C, Inactive lever: Treatment F3,54=19.68, P < 0.01; Treatment × Sex F3,54=2.788, P < 0.05; Sex F1,18=0.166, NS). Post hoc comparisons revealed that SCH23390 significantly reduced inactive lever responding in males at all doses tested (0.003, 0.01, and 0.03 mg/kg), whereas in females, a reduction was observed only at the highest dose (0.03 mg/kg) (Fig 3C).

**Fig. 3.**
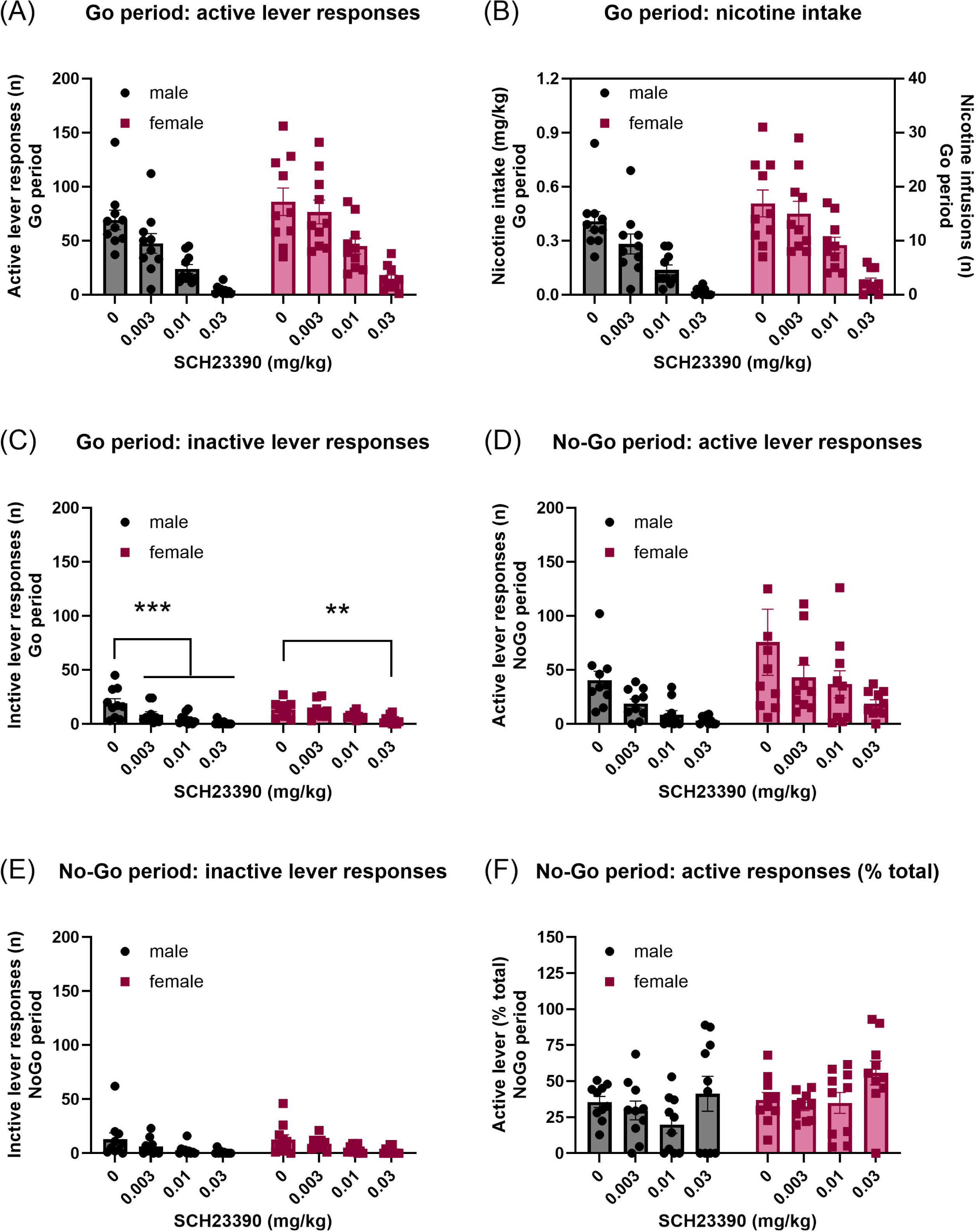
Effects of SCH23390 on performance in the nicotine Go/No-Go task. Effects of the dopamine D1 receptor antagonist SCH23390 on (A) active lever responses, (B) nicotine intake, and (C) inactive lever responses during the Go periods. SCH23390 reduced active lever responding and nicotine intake during Go periods. SCH23390 also reduced inactive lever responding, with males showing reduced responding at all doses and females showing reduced responding only at the highest dose. Effects of SCH23390 on (D) active lever responses, (E) inactive lever responses, and (F) the percentage of active lever responses during the No-Go periods. SCH23390 reduced responding during No-Go periods and affected the percentage of active lever responses. Males, N=10; females, N=10. ** p < 0.01, *** p < 0.001. Data are expressed as mean ± SEM.

#### No-Go period

During the No-Go period, SCH23390 treatment decreased both active and inactive lever responses in both male and female rats (Fig. 3D, Active lever: Treatment F3,54=7.573, P < 0.01; Treatment × Sex F3,54=0.318, NS; Sex F1,18=3.811, NS; Fig. 3E, Inactive lever: Treatment F3,54=8.185, P < 0.01; Treatment × Sex F3,54=0.138, NS; Sex F1,18=0.159, NS). Similarly, SCH23390 significantly affected the percentage of active lever responses during the No-Go period (Fig. 3F, Treatment F3,54=4.19, P < 0.01; Treatment × Sex F3,54=0.657, NS; Sex F1,18=1.807, NS).

### 3.4. Go/No-Go task: A77636 treatment

#### Go period

A77636 treatment reduced both active lever responses and nicotine intake during the Go period in both males and females (Fig. 4A, Active lever: Treatment F3,51=6.106, P < 0.001; Treatment × Sex F3,51=1.041, NS; Sex F1,17=0.745, NS; Fig. 4B, Nicotine intake: Treatment F3,51=7.507, P < 0.001; Treatment × Sex F3,51=1.666, NS; Sex F1,17=0.521, NS). In contrast, A77636 treatment did not affect inactive lever responses (Fig. 4C, Treatment F3,51=1.821, NS; Treatment × Sex F3,51=0.42, NS; Sex F1,17=0.162, NS).

**Fig. 4.**
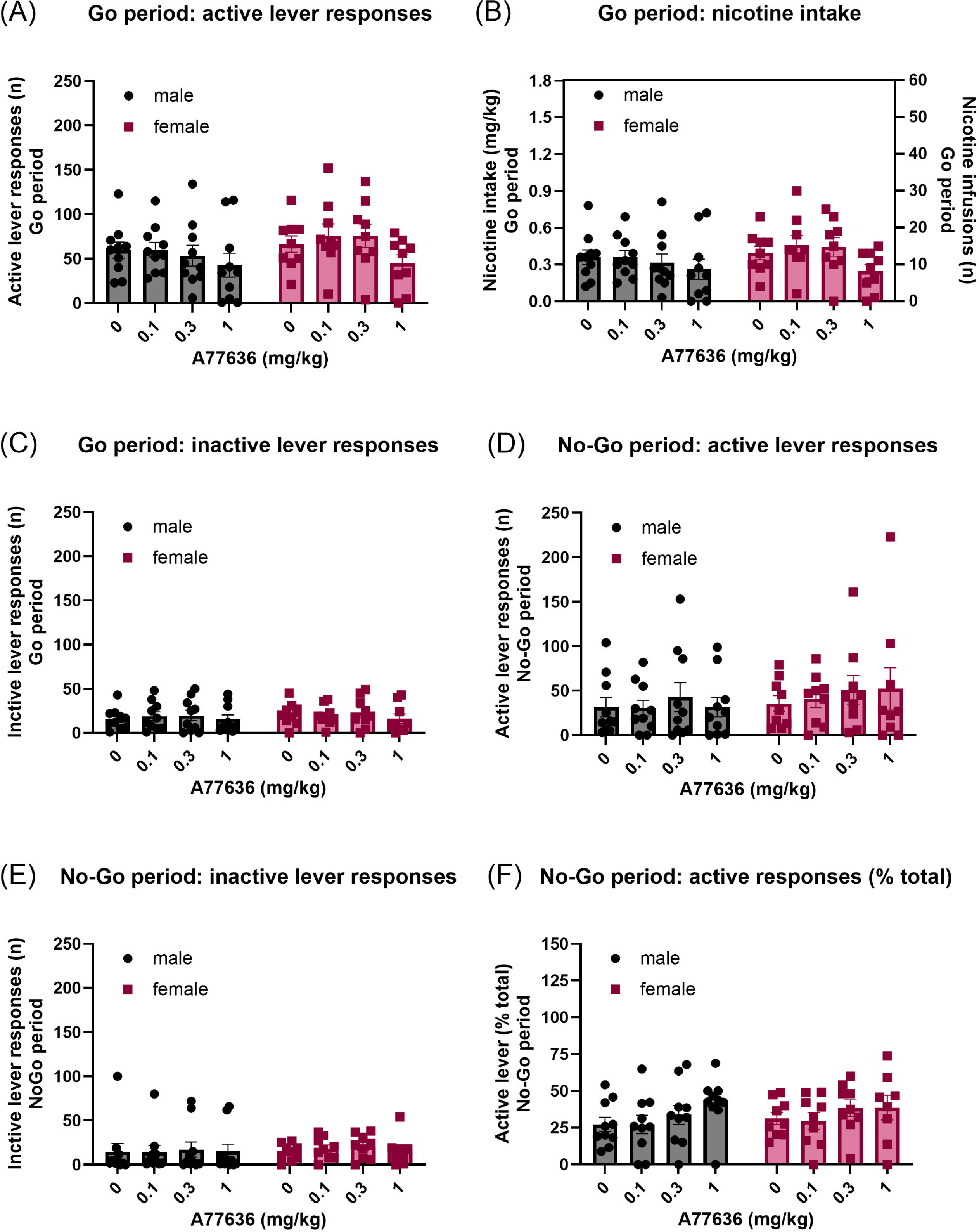
Effects of A77636 on performance in the nicotine Go/No-Go task. Effects of the dopamine D1 receptor agonist A77636 on (A) active lever responses, (B) nicotine intake, and (C) inactive lever responses during the Go periods. A77636 reduced active lever responding and nicotine intake during Go periods but did not significantly alter inactive lever responding. Effects of A77636 on (D) active lever responses, (E) inactive lever responses, and (F) the percentage of active lever responses during the No-Go periods. A77636 did not significantly affect No-Go responding or the percentage of active lever responses during No-Go periods, indicating that D1 receptor stimulation reduced nicotine-motivated responding without altering the primary measure of impulsive action. Males, N=10; females, N=9. Data are expressed as mean ± SEM.

#### No-Go period

During the No-Go period, A77636 treatment did not affect either the active or inactive lever responses (Fig. 4D, Active lever: Treatment F3,51=0.876, NS; Treatment × Sex F3,51=0.288, NS; Sex F1,17=0.449, NS; Fig. 4E, Inactive lever: Treatment F3,51=0.579, NS; Treatment × Sex F3,51=0.216, NS; Sex F1,17=0.042, NS). Similarly, A77636 treatment did not affect the percentage of active lever responses during the No-Go period (Fig. 4F, treatment F3,51=2.599, NS; Treatment × Sex F3,51=0.258, NS; Sex F1,17=0.12, NS).

### 3.5. Go/No-Go task: Saline and nicotine self-administration

#### Go period

During the Go period, active and inactive lever responses, as well as the number of infusions earned, did not differ between the Nicotine and Saline groups in either males or females (Fig. 5A, Active lever: Group F1,17=3.666, NS; Group × Sex F1,17=0.28, NS; Sex F1,17=0, NS; Fig. 5B, Infusions: Group F1,17=4.008, NS; Group × Sex F1,17=0.533, NS; Sex F1,17=0.002, NS; Fig. 5C, Inactive lever Group F1,17=6.19, NS; Group × Sex F1,17=0.095, NS; Sex F1,17=0.117, NS).

**Fig. 5.**
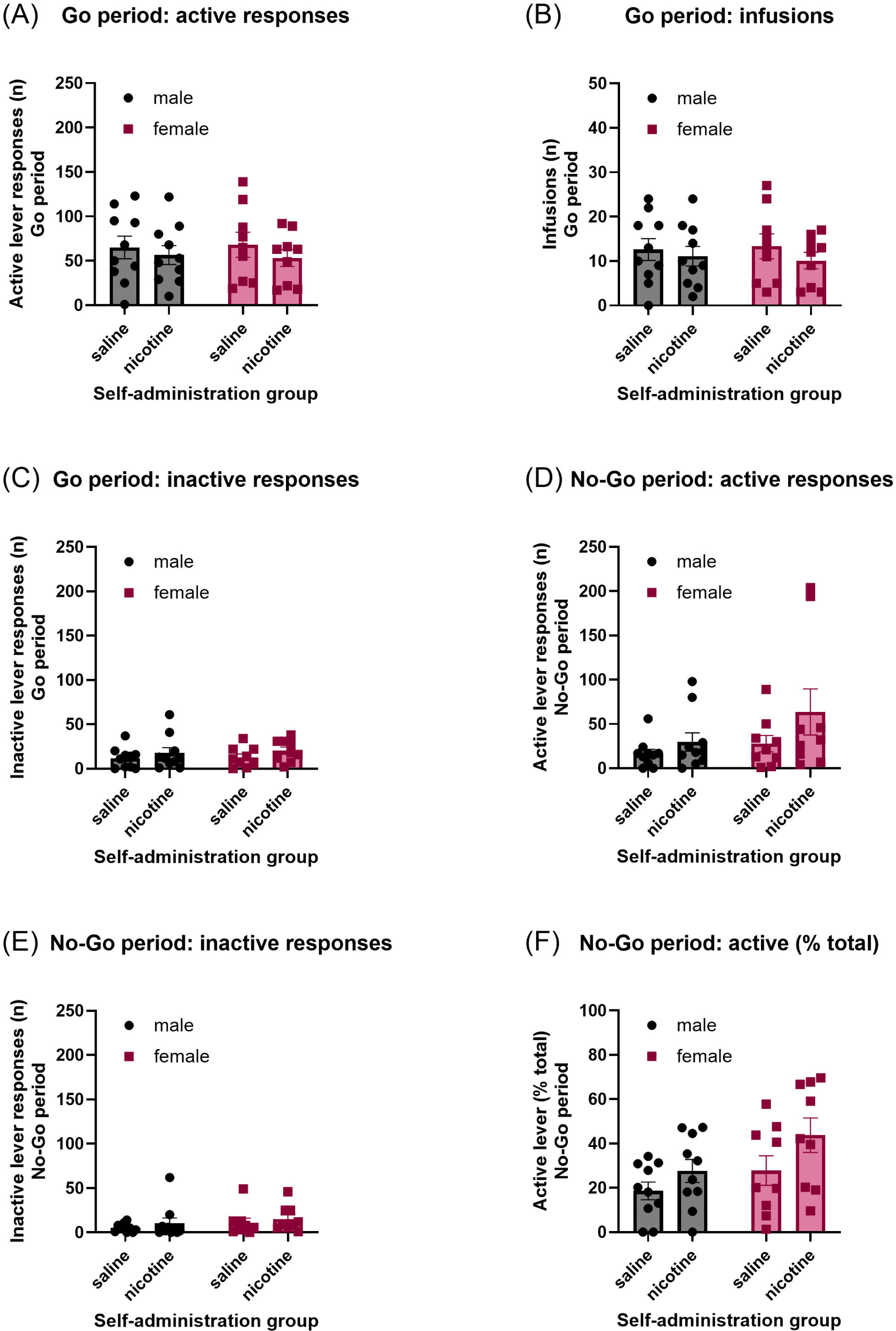
Comparison of responding in rats self-administering nicotine versus saline in the Go/No-Go task. (A) Active lever responses, (B) total infusions, and (C) inactive lever responses during Go periods in rats self-administering nicotine or saline. Go-period responding and infusions did not differ significantly between nicotine and saline self-administration conditions. (D) Active lever responses, (E) inactive lever responses, and (F) the percentage of active lever responses during No-Go periods. Rats self-administering nicotine showed higher active lever responding and a higher percentage of active lever responses during No-Go periods than rats self-administering saline, indicating that nicotine increased impulsive action without increasing overall Go-period operant responding. Males, N=10; females, N=9. Data are expressed as mean ± SEM.

#### No-Go period

During the No-Go period, rats in the Nicotine Group exhibited more active lever responses than those in the Saline Group, with no significant sex differences (Fig 5D, Group F1,17=5.763, P < 0.05; Group × Sex F1,17=1.218, NS; Sex F1,17=1.657, NS). In contrast, inactive lever responses did not differ between the Nicotine and Saline groups (Fig 5E, Group F1,17=2.054, NS; Group × Sex F1,17=0.03, NS; Sex F1,17=0.372, NS). The percentage of active lever responses during the No-Go period was higher in the Nicotine Group compared with the Saline Group, with no sex differences observed (Fig 5F, Group F1,17=17.583, P < 0.001; Group × Sex F1,17=1.373, NS; Sex F1,17=2.585, NS).

### 3.6. Go/No-Go task: mecamylamine treatment

#### Go period

Mecamylamine treatment decreased both active lever responses and nicotine intake during the Go period, with significant sex differences observed (Fig. 6A, Active lever: Treatment F1,17=8.455, P = 0.01; Treatment × Sex F1,17=0.436, NS; Sex F1,17=5.326, P < 0.05; Fig. 6B, Nicotine intake: Treatment F1,17=10.264, P < 0.01; Treatment × Sex F1,17=1.162, NS; Sex F1,17=5.002, P < 0.05). Mecamylamine treatment also decreased inactive lever responses during the Go period, with no sex differences observed (Fig. 6C, Inactive lever: Treatment F1,17=5.121, P < 0.05; Treatment × Sex F1,17=0.712, NS; Sex F1,17=1.301, NS).

**Fig. 6.**
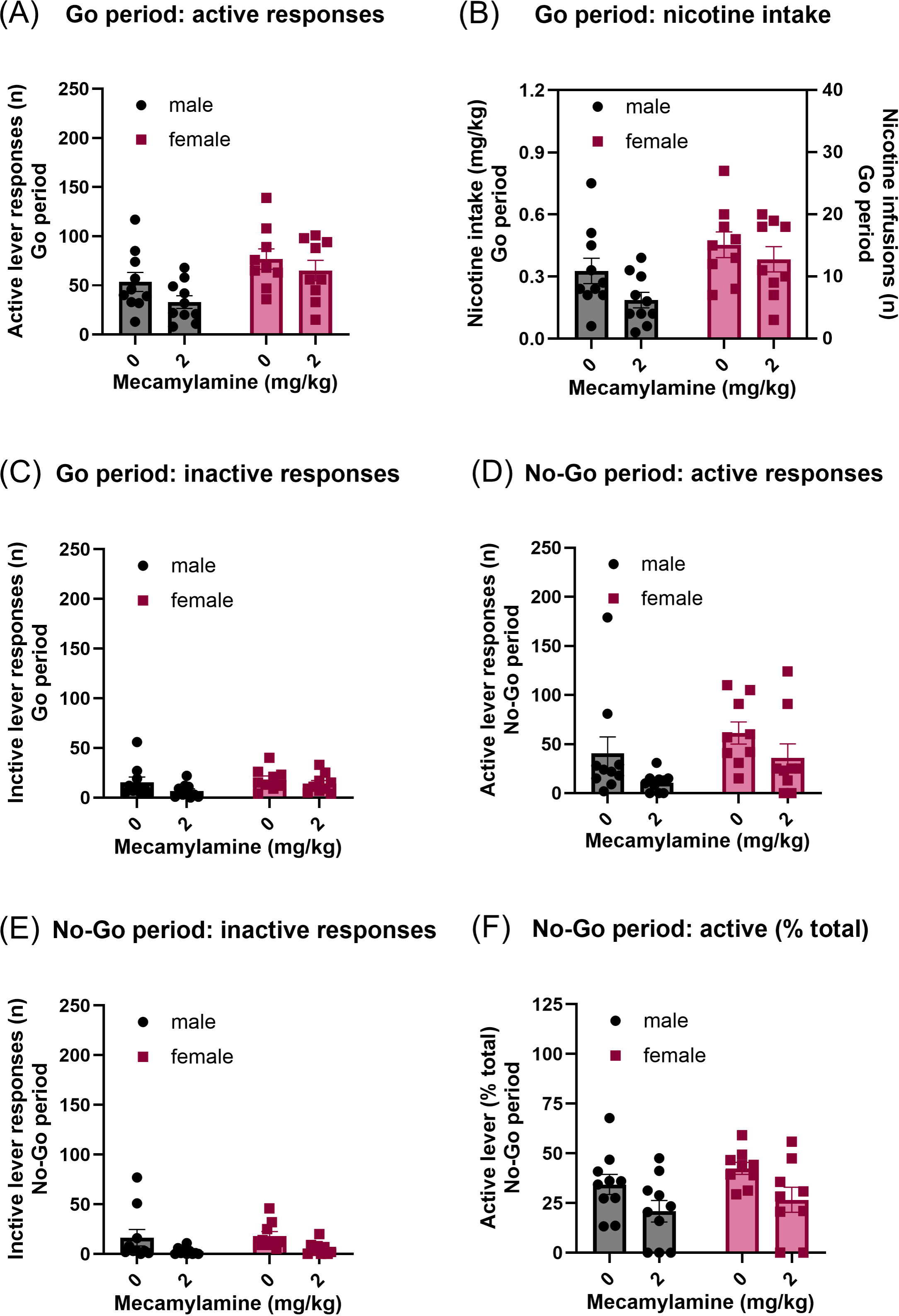
Effects of mecamylamine on performance in the nicotine Go/No-Go task. Effects of the non-selective nAChR receptor antagonist mecamylamine on (A) active lever responses, (B) nicotine intake, and (C) inactive lever responses during the Go periods. Mecamylamine reduced active lever responding, nicotine intake, and inactive lever responding during Go periods. Effects of mecamylamine on (D) active lever responses, (E) inactive lever responses, and (F) the percentage of active lever responses during the No-Go periods. Mecamylamine reduced both active and inactive lever responding during No-Go periods and decreased the percentage of active lever responses, consistent with reduced nicotine-related impulsive action. Males, N=10; females, N=9. Data are expressed as mean ± SEM.

#### No-Go period

During the No-Go period, mecamylamine treatment decreased both active and inactive lever responses in both males and females (Fig. 6D, Active lever: Treatment F1,17=10.602, P < 0.01; Treatment × Sex F1,17=0.087, NS; Sex F1,17=2.264, NS; Fig. 6E, Inactive lever: Treatment F1,17=7.798, P < 0.05; Treatment × Sex F1,17=0.009, NS; Sex F1,17=0.187, NS). Mecamylamine treatment also decreased the percentage of active lever responses during the No-Go period (Fig. 6F, Treatment F1,17=13.158, P < 0.01; Treatment × Sex F1,17=0.085, NS; Sex F1,17=1.328, NS).

## DISCUSSION

The present study examined the role of dopamine D1 receptor signaling in nicotine-motivated responding and impulsive action using a Go/No-Go intravenous self-administration procedure in male and female rats. During Go/No-Go training, active lever responding and nicotine intake increased during the initial Go periods and then stabilized, while the percentage of active lever responses during No-Go periods progressively decreased before reaching a stable level. Together these findings demonstrate that rats learned to discriminate between periods of nicotine availability and unavailability. A significant main effect of sex was also observed during training, with females showing a higher percentage of active lever responses during No-Go periods than males, indicating a baseline sex difference in inhibitory control. Following training, the D1 receptor antagonist SCH23390 reduced nicotine intake and active lever responding during Go periods. SCH23390 also affected the percentage of active lever responses during No-Go periods. However, this effect was accompanied by reduced inactive lever responding and is therefore difficult to attribute specifically to a change in nicotine-induced impulsive action rather than to a general reduction in operant output. In contrast, the D1 receptor agonist A77636 reduced nicotine-motivated responding during Go periods without affecting impulsive action during No-Go periods, indicating that D1 receptor signaling contributes primarily to nicotine-motivated responding rather than to nicotine-induced impulsive action. Rats displayed significantly greater impulsive action during periods of active nicotine self-administration compared with saline self-administration, confirming that the elevated percentage of active lever responses during No-Go periods reflected an acute pharmacological effect of nicotine rather than a fixed deficit that persists in the absence of the drug. Mecamylamine reduced nicotine-motivated responding during Go periods and decreased the percentage of active lever responses during No-Go periods, indicating that blockade of nAChRs reduced impulsive action. Together, these findings suggest that nicotine-induced impulsive action in this task is mediated primarily through nicotinic acetylcholine receptor signaling, whereas dopamine D1 receptor signaling contributes mainly to nicotine-motivated responding.

### Active nicotine self-administration increases impulsive action

Throughout this study, impulsive action was indexed by the percentage of active lever responses emitted during No-Go periods relative to total active lever responses. Because this measure expresses No-Go responding as a proportion of total active responding, it helps account for differences in overall response output and provides a response-allocation measure of inhibitory control. The comparison between nicotine and saline self-administering rats provided important evidence for the pharmacological specificity of the No-Go measure in this task. During Go periods, active lever responding and the number of infusions earned did not differ between the nicotine and saline groups. In contrast, rats self-administering nicotine showed significantly higher active lever responding and a higher percentage of active lever responses during No-Go periods compared with saline controls, indicating that the elevated No-Go responding in nicotine rats reflected a specific effect of nicotine on impulsive action. This finding is consistent with evidence that nicotine impairs response inhibition (Kolokotroni et al. 2011; 2012). This dissociation is informative because responding during Go periods was maintained at comparable levels in both conditions, likely through the conditioned reinforcing properties of the nicotine-paired cue, the two groups were matched for operant output and number of reinforcers earned. The selective elevation of No-Go responding in the nicotine group therefore cannot be explained by differences in overall responding or cue-maintained behavior and instead reflects the pharmacological presence of nicotine. This is consistent with the well-characterized pattern of responding when nicotine is withheld during self-administration, in which lever responding is initially maintained, or transiently increased, before declining gradually across sessions rather than ceasing abruptly (Liu et al. 2006; Macnamara et al. 2016; Shram et al. 2008). Within a single saline-substitution session, responding therefore remains near nicotine-maintained levels, so that the comparable Go-period responding in the two groups reflects this persistence of cue- and history-maintained behavior, whereas the loss of elevated No-Go responding reflects the pharmacological absence of nicotine. Together, this pattern dissociates the pharmacological contribution of nicotine to impulsive action from its effects on operant responding. This indicates that the elevated impulsive action observed during nicotine self-administration depends on the acute presence of the drug rather than reflecting a persistent deficit in inhibitory control that would remain when nicotine is unavailable. No sex differences were observed in the saline versus nicotine comparison, suggesting that the elevation in impulsive action by nicotine was similar across sexes.

### Nicotinic Acetylcholine receptor blockade reduces nicotine-induced impulsive action

The non-selective nAChR antagonist mecamylamine reduced nicotine intake and active lever responding during Go periods and decreased the percentage of active lever responses during No-Go periods. Because the percentage measure is independent of the overall rate of responding, its reduction indicates a shift in response allocation away from the nicotine-associated lever, rather than a general suppression of behavior. Mecamylamine therefore reduced impulsive action, supporting a major role for nAChR activation in the elevated impulsive action observed during nicotine self-administration. This is consistent with our previous finding that nAChR blockade reduced impulsive action in the same Go/No-Go nicotine self-administration procedure (Chellian et al. 2026). This interpretation is also consistent with evidence that acute nicotine increases behavioral disinhibition in rats and that this effect is blocked by mecamylamine (Kolokotroni et al. 2011). In the 5-CSRTT, nicotine-induced increases in premature responding are blocked by mecamylamine and by the α4-containing nAChR antagonist dihydro-β-erythroidine, but not by the α7 antagonist methyllycaconitine, indicating that α4-containing nAChRs mediate the effect (Blondel et al. 2000; Jepsen et al. 2014; Papke et al. 2008). The involvement of nAChRs in impulsive action is further supported by evidence that mecamylamine reduces premature responding in the 5-CSRTT (Balachandran et al. 2018). Significant main effects of sex were observed for active lever responding and nicotine intake during the Go period, with males showing higher overall responding than females; however, there were no Treatment × Sex interactions for any measure, indicating that the effects of mecamylamine did not differ between sexes. Although mecamylamine was not tested in saline self-administering rats in the present study, earlier work indicates that it does not affect impulsive responding in drug-naive animals in symmetrically reinforced Go/No-Go, delayed-reward, or 5-CSRTT paradigms (Blondel et al. 2000; Kolokotroni et al. 2011). This suggests that the effect of mecamylamine observed here reflects blockade of nicotine-driven impulsivity rather than a nonspecific suppression of responding.

### Dopamine D1 receptor blockade reduces nicotine-motivated responding without a selective effect on impulsive action

SCH23390 dose-dependently reduced nicotine intake and active lever responding during Go periods, consistent with the well-established role of D1 receptor signaling in nicotine reinforcement. These findings are in agreement with prior work showing that SCH23390 reduces nicotine self-administration in rats (Chellian et al. 2022; Corrigall and Coen 1991). Significant main effects of sex were observed for active lever responding and nicotine intake during the Go period, with females showing higher overall responding than males. However, the Treatment × Sex interaction was not significant, indicating that SCH23390 reduced nicotine-motivated responding similarly in both sexes.

Importantly, SCH23390 also affected the percentage of active lever responses during No-Go periods. However, this effect was statistically a main effect of treatment without a significant interaction, and a clear dose-dependent direction could not be established. Moreover, SCH23390 concurrently reduced active lever responding, inactive lever responding, and nicotine intake during both Go and No-Go periods, indicating a broad reduction in operant output rather than a selective effect on inhibitory control. This interpretation is reinforced by our previous finding that, at the same doses and pretreatment interval used here, SCH23390 decreased nicotine intake, food-maintained responding, and locomotor activity (Chellian et al. 2022). Because SCH23390 suppressed responding across reinforcers and reduced locomotor activity, its effect on the No-Go measure in the present task cannot be dissociated from a general reduction in motor and operant output and should not be interpreted as a specific improvement in inhibitory control. This limitation is specific to the present task rather than to D1 receptors as such. In the 5-CSRTT, where premature responding can be separated from overall response rate, SCH23390 reduces premature responding and attenuates nicotine-induced impulsivity (Balachandran et al. 2018), indicating that D1 receptor signaling can contribute to impulsive action under conditions that permit this dissociation. The Go/No-Go self-administration procedure used here, in which SCH23390 also reduced nicotine-motivated responding and locomotor output, does not allow the contribution of D1 receptor blockade to inhibitory control to be separated from its effect on overall responding.

### Dopamine D1 receptor stimulation reduces nicotine-motivated responding without affecting impulsive action

The D1 receptor agonist A77636 reduced active lever responding and nicotine intake during Go periods without affecting responding during No-Go periods. A77636 also did not affect inactive lever responding. This selective reduction in nicotine-motivated responding, without a corresponding change in impulsive action, indicates that D1 receptor stimulation reduced the motivation to obtain nicotine while leaving inhibitory control intact. The reduction in Go-period responding by A77636 is consistent with previous reports that D1-like receptor agonists can suppress operant responding for nicotine and food (Chellian et al. 2022). Importantly, at the doses and pretreatment interval used here, A77636 reduces nicotine intake without reducing locomotor activity (Chellian et al. 2022), indicating that this effect on nicotine-motivated responding is not secondary to motor impairment. The contrast between the two D1 compounds is informative. SCH23390 reduced nicotine-motivated responding together with a broad reduction in operant and motor output, whereas A77636 reduced nicotine-motivated responding selectively, without affecting impulsive action or locomotor activity. The selective effect of A77636 indicates that a reduction in nicotine-motivated responding is not sufficient to alter impulsive action. Considered alongside the reduction of impulsive action by nAChR receptor blockade, this pattern indicates that nicotine-induced impulsive action in this task is mediated primarily through nAChR signaling, whereas D1 receptor signaling contributes mainly to nicotine-motivated responding. This dissociation has potential translational relevance, as it suggests that D1 receptor stimulation can reduce nicotine-motivated behavior without worsening impulsive action in a way that could increase relapse vulnerability. No sex differences were observed for any measure during A77636 treatment, indicating that D1 receptor stimulation affected nicotine-motivated responding similarly in males and females.

### Sex differences in impulsive action

The present study also revealed sex-related differences in impulsive action during Go/No-Go training. Females showed a higher percentage of active lever responses during No-Go periods than males, suggesting greater baseline impulsive action. This finding is consistent with clinical evidence that impulsivity is an important vulnerability factor for substance use disorders, although sex differences in impulsivity are complex and depend on the specific task and population studied (Verdejo-García et al. 2008). Sex differences in Go/No-Go performance are not uniform across studies. For example, some human studies report better No-Go inhibition in women than men, indicating that sex effects may depend on task parameters, smoking status, and clinical characteristics (Sjoberg and Cole 2018). Preclinical studies also indicate that sex differences in impulsive action depend on the behavioral model and reinforcer. In general, males often show greater impulsive action in some food-maintained tasks, whereas females may show greater impulsivity under drug-related conditions or in tasks that involve drug-associated motivation (Hamilton et al. 2015; Weafer and de Wit 2014). Thus, the higher No-Go responding observed in females in the present nicotine self-administration task may reflect sex differences in the interaction between nicotine motivation and inhibitory control, rather than a generalized increase in impulsivity. Importantly, although sex differences were observed during training and in some measures of operant responding after SCH23390 and mecamylamine treatment, there were no Treatment × Sex interactions for the primary measure of impulsive action after SCH23390, A77636, or mecamylamine treatment. These findings suggest that baseline inhibitory control differed between males and females, but pharmacological modulation of nicotine-related impulsive action was similar across sexes.

### Limitations

Several limitations should be considered when interpreting the present findings. First, SCH23390 reduced operant responding across both Go and No-Go periods, reduced inactive lever responding, and has previously been shown to reduce locomotor activity and food-maintained responding at the doses and pretreatment interval used here (Chellian et al. 2022). Therefore, the effects of SCH23390 on No-Go responding should be interpreted in the context of a broader reduction in operant and motor output, which limits conclusions about the specific contribution of D1 receptor blockade to inhibitory control. Second, the study used systemic drug administration, which does not identify the specific brain regions or circuits through which D1 receptor signaling modulated nicotine-motivated responding and impulsive action. Future studies using site-specific pharmacological manipulations or circuit-based approaches will be needed to determine the relative contributions of the PFC, NAc, and dorsal striatum. Third, the present study examined only D1-selective compounds, and the contribution of D2-like receptors to nicotine-induced impulsive action was not assessed. This is relevant because the D2 receptor antagonist eticlopride attenuates nicotine-induced impulsivity in the 5-CSRTT (Balachandran et al. 2018), indicating that D2 receptor signaling may also contribute to nicotine-related impulsive action. Future work using D2-selective agonists and antagonists will be needed to define the relative contributions of D1 and D2 receptors. Finally, although sex was included as a biological variable, the study was not designed to determine the role of the estrous cycle in nicotine-related impulsive action. However, previous studies found no evidence that the estrous cycle affects nicotine reward or intake (Donny et al. 2000; Torres et al. 2009).

## Conclusion

In conclusion, the present study demonstrates that nicotine self-administration increases impulsive action, and that nicotine-motivated responding and impulsive action can be pharmacologically dissociated. The D1 receptor agonist A77636 selectively reduced nicotine-motivated responding without affecting impulsive action, indicating that D1 receptor signaling contributes mainly to the motivation to obtain nicotine. The D1 receptor antagonist SCH23390 reduced nicotine-motivated responding but also broadly suppressed operant and motor output, so its effect on the impulsivity measure could not be attributed specifically to a change in inhibitory control. In contrast, the non-selective nAChR antagonist mecamylamine reduced impulsive action, identifying nAChR receptor signaling as a primary mechanism underlying the elevated impulsive action observed during nicotine self-administration. The finding that rats showed greater impulsive action during nicotine self-administration than during saline self-administration indicates that elevated impulsive action depends on the acute presence of nicotine rather than a persistent deficit in inhibitory control. Sex differences in baseline impulsive action were observed, with females showing greater No-Go responding during training. Together, these findings indicate that nicotine-induced impulsive action is mediated primarily through nAChR receptor signaling, whereas D1 receptor signaling contributes mainly to nicotine-motivated responding, and suggest that selective D1 receptor stimulation warrants further investigation as a strategy for reducing nicotine-motivated behavior without affecting impulsive action.

## CRediT authorship contribution statement

**Adriaan W. Bruijnzeel:** Conceptualization, Supervision, Formal analysis, Writing – Original Draft, Visualization, Project administration, Funding acquisition.

**Guido Huisman:** Investigation, Project administration, Writing – review & editing.

**Lara Caglayan:** Investigation, Project administration, Writing – review & editing.

**Ranjithkumar Chellian:** Conceptualization, Formal analysis, Investigation, Writing – review & editing, Visualization.

### Data Integrity and Sponsor Role

The authors confirm that they have had full access to all the data in the study and take responsibility for the integrity of the data and the accuracy of the data analysis. No sponsor was involved in the study design, data collection, or writing of the manuscript.

### Design and Analysis Transparency

The datasets generated and analyzed during the current study are available from the corresponding author on request.

