## Supplemental Table S1 and Figure S1 for "Dissociable roles of dopamine D1 and nicotinic receptors in nicotine-motivated responding and impulsive action in a Go/No-Go self-administration task"

**Table S1. Animal numbers and exclusions across experimental phases.**

| **Study phase** | **Exclusion during phase** | **Males remaining** | **Females remaining** | **N to report** |
| --- | --- | --- | --- | --- |
| Start of study | None | 14 | 18 | Males N=14; females N=18 |
| Acquisition of nicotine self-administration | 1 male and 4 females failed the Brevital catheter patency test | 13 | 14 | Males N=13; females N=14 |
| Go/No-Go training | 3 males and 3 females failed the Brevital test | 10 | 11 | Males N=10; females N=11 |
| SCH23390 treatment phase | 1 additional female failed the Brevital test | 10 | 10 | Males N=10; females N=10 |
| A77636 treatment phase | 1 additional female failed the Brevital test | 10 | 9 | Males N=10; females N=9 |
| Saline versus nicotine comparison | No additional exclusions | 10 | 9 | Males N=10; females N=9 |
| Mecamylamine treatment phase | No additional exclusions | 10 | 9 | Males N=10; females N=9 |

**Figures**

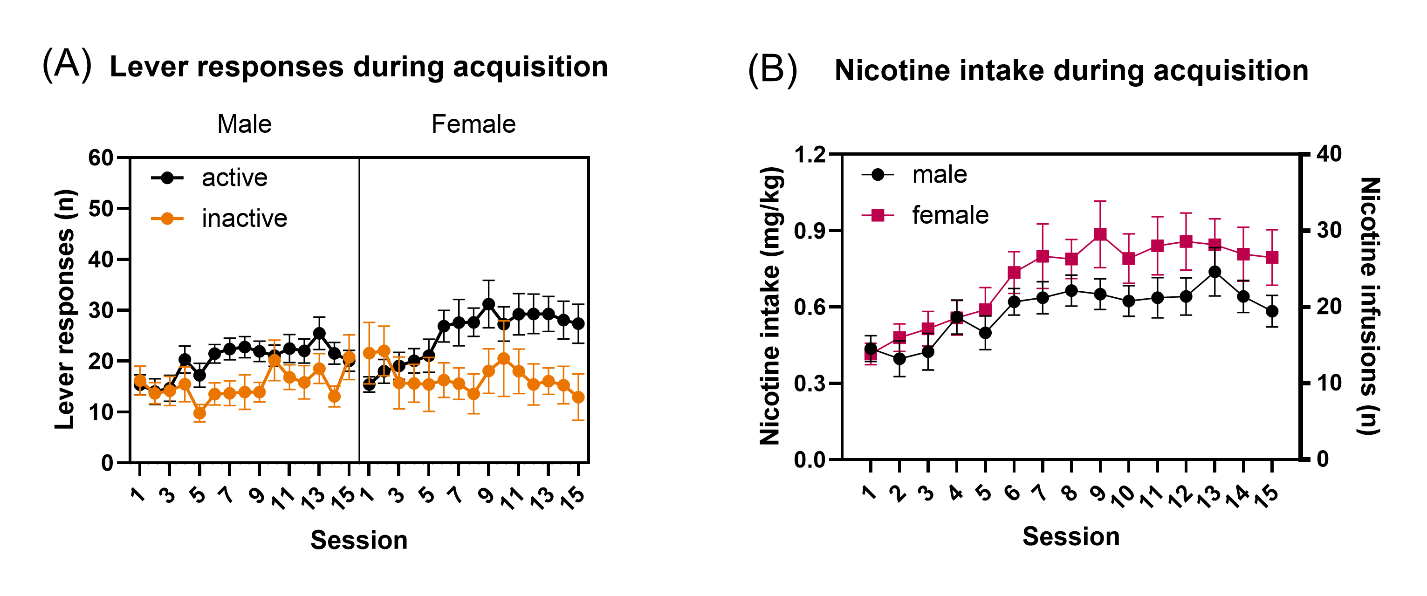
**Fig. S1. Acquisition of nicotine self-administration.** (A) Active and inactive lever responses during the 15 acquisition sessions in male and female rats. Rats responded more on the active lever than on the inactive lever, indicating acquisition of nicotine self-administration. Active lever responding increased during the initial sessions and then stabilized, whereas inactive lever responding remained relatively low across sessions. (B) Nicotine intake and number of nicotine infusions earned during acquisition. Nicotine intake increased during the initial sessions and then stabilized in both males and females. Males, N=13; females, N=14. Data are expressed as mean ± SEM.
